# Short-term block of CRH receptor in adults mitigates age-related memory impairments provoked by early-life adversity

**DOI:** 10.1101/714451

**Authors:** Annabel K. Short, Pamela M. Maras, Aidan L. Pham, Autumn S. Ivy, Tallie Z. Baram

## Abstract

In humans, early-life adversity (ELA) is associated with impairments in learning and memory that may emerge later in life. In rodent models, ELA directly impacts hippocampal neuron structure and connectivity with progressive deficits in long-term potentiation and spatial memory function. Previous work has demonstrated that augmented release and actions of the stress-activated neuropeptide, CRH, contribute to the deleterious effects of ELA on hippocampal structure and memory-function. Early-life adversity increases CRH production and levels, and blocking CRH receptor type 1 (CRHR1) within the hippocampus immediately following adversity prevented the memory and LTP problems caused by ELA. Here we queried if blocking CRHR1 during adulthood ameliorates the adverse impact of ELA on memory in middle age. Blocking CRHR1 for a week in two month old male rats prevented ELA-induced deficits in object recognition memory that emerge during middle age. The intervention failed to mitigate the reduction of spatial memory at 4 and 8 months, but restored hippocampus-dependent location memory in ELA-experiencing rats during middle age (12 months of age).

Notably, neither ELA nor blocking CRHR1 influenced anxiety- or depression-related behaviors These findings suggest a sensitive period during which interventions can fully prevent long-lasting effects of ELA, yet indicate that interventions later in life offer significant benefits.

## Introduction

Age-related memory loss has a significant impact on an individual’s quality of life in addition to global economic burden (Prince, Guerchet, & Prina, 2013; Prince et al., 2015). Predisposition to cognitive disorders throughout life is established through an interplay of inherited and environmental factors (Caspi et al., 2003; Klengel & Binder, 2015). Brain development during the early postnatal period is particularly susceptible to environmental influences (Brown, Susser, Lin, Neugebauer, & Gorman, 1995; Chen & Baram, 2016; Eriksson, Räikkönen, & Eriksson, 2014; Lupien, McEwen, Gunnar, & Heim, 2009; Novick et al., 2018; Raymond, Marin, Majeur, & Lupien, 2018). In humans, an impoverished environment during childhood is associated with impaired cognition later in life, and an increased risk of dementia (Kaplan et al., 2001; Nelson et al., 2007). Stress early in life can lead to delayed, progressive impairments of hippocampal function (Brunson et al., 2005). These enduring memory deficits are likely due to a cascade of cellular and molecular mechanisms that ultimately result in changes to learning and memory circuits (Schulmann et al.; Singh-Taylor, Korosi, Molet, Gunn, & Baram, 2015; Singh-Taylor et al., 2018).

The hippocampus is particularly vulnerable to adverse experiences early in life. There is evidence of reduced hippocampal volume in children raised in orphanages (Hodel et al., 2015) and rodents exposed to early-life stress (Brunson et al., 2005; Ivy et al., 2010; Molet, Maras, et al., 2016). Investigations in rodents suggest decreases in hippocampal volume following early life adversity are a result of reduced dendritic arborization (Molet, Maras, et al., 2016). Reduction in hippocampal dendrites is associated with chronic increases in corticotropin releasing hormone (CRH) (Chen et al., 2013; Maras & Baram, 2012). CRH is expressed in the hippocampus within a subpopulation of interneurons (Chen et al., 2004; Chen, Bender, Frotscher, & Baram, 2001; Gunn, Sanchez, Lynch, Baram, & Chen, 2019; Hooper, Fuller, & Maguire, 2018; Hooper & Maguire, 2016; Yan, Toth, Schultz, Ribak, & Baram, 1998) and binds to corticotropin releasing hormone receptor type −1 (CRHR1) receptors on pyramidal cells, resulting in neuronal activation immediately following stress (Chen, Brunson, Müller, Cariaga, & Baram, 2000). Sustained increases in CRH binding to CRHR1 in the hippocampus results in reduced dendritic spines and synapse integrity via actin remodeling (Chen, Dube, Rice, & Baram, 2008; Chen et al., 2013, 2010), which results in deficits in learning and memory (Kim & Diamond, 2002; McEwen, 1999).

In a rodent model of early life adversity, which simulates an impoverished environment through Limited Bedding and Nesting (LBN) and induces disrupted patterns of maternal care, there is elevated CRH in the hippocampus, both immediately following LBN and in adult animals (Fenoglio, Brunson, & Baram, 2006; Ivy et al., 2010; Maras & Baram, 2012). As adults, rodents who experienced LBN early in life have significant impairments in learning and memory (Bath, Manzano-Nieves, & Goodwill, 2016; Brunson et al., 2005; Molet, Maras, et al., 2016; Naninck et al., 2015; Rice, Sandman, Lenjavi, & Baram, 2008; Walker et al., 2017), which worsen with age (Brunson, Eghbal-Ahmadi, Bender, Chen, & Baram, 2001) and following a second stress (Molet, Maras, et al., 2016). These impairments in learning and memory were replicated by infusing CRH directly into the brains of immature rats while controlling the levels of circulating glucocorticoids (Brunson et al., 2001, 2005). Conversely, when a CRHR1 antagonist is administered during the sensitive period of hippocampal development, deficits in learning and memory following LBN are prevented (Ivy et al., 2010). This suggests a vital contribution of CRH in the progressive deficits in learning and memory observed following early life adversity.

Understanding how early experiences alter brain circuits long term, and the effectiveness of interventions across the lifespan, is important for identifying treatment options and outcomes. The goal of the present study was to identify if interventions outside of the sensitive period, when memory impairments are beginning to emerge, could also prevent the progression of memory loss.

## Methods

### Animals

Subjects were male rats born to timed-pregnant Sprague-Dawley rat dams maintained on 12 h light/ dark cycles with *ad libitum* access to chow and water. On P2, pups from multiple litters were gathered, and 12 (6 males; 6 females) were assigned at random to each dam, to obviate potential genetic and litter size confounders. After weaning, males were housed three per cage. All experiments were performed in accordance with National Institutes of Health guidelines and were approved by the University of California, Irvine, Animal Care and Use Committee.

### The Early-Life Adversity Paradigm

The adversity paradigm/ Limited bedding nesting paradigm (LBN) consisted of limiting nesting and bedding materials in cages between P2-P9 (Molet, Maras, Avishai-Eliner, & Baram, 2014). For the LBN group, a plastic-coated aluminum mesh platform was placed ∼2.5cm above the floor of a standard cage. Cobb bedding was reduced to cover cage floor in a single layer, and one-half of a single paper towel was provided for nesting material on the platform. Control dams and litters resided in standard cages containing ample cob bedding and one whole paper towel for nesting. This paradigm causes maternal care to be fragmented and unpredictable, provoking chronic stress in the pups (Brunson et al., 2005; Ivy, Brunson, Sandman, & Baram, 2008; Molet et al., 2014; Molet, Maras, et al., 2016). Control and experimental cages were undisturbed during P2–P9, housed in temperature-controlled rooms (22°C). On P10, experimental groups were transferred to standard cages, where maternal behavior normalized within hours (Ivy et al., 2008; Molet et al., 2014). Rats were weaned on P21-22, and then group housed. Testing for emotional and cognitive disorders were performed at four, eight and again at 12 months. One animal in the LBN antagonist group died prior to FST and EPM testing at 12 months of age and therefore is missing from these analyses.

### Intracerebroventricular (ICV) Administration of CRFR1 antagonist

The selective CRFR1 blocker, NBI30775(3-[6-(dimethylamino)-4-methyl-pyrid-3-yl]-2,5-dimethyl-N,N-dipropyl-pyrazolo[2,3-a]pyrimidin-7-amine) or vehicle was chronically infused intra-cerebroventricular (ICV) via osmotic minipumps to control and LBN male rats for one week, at 2 months of age (Ivy et al., 2010). Briefly, NBI30775 was dissolved in warm distilled water with 5.5% cremophor EL and the pH was adjusted; vehicle solution was made using the same pH-adjusted distilled water and 5.5% cremophor without the compound. Drug solution (11 mg/ml) or vehicle was loaded into osmotic Alzet minipumps (model 2001, Alzet Corp., Cupertino, CA), calibrated to release 0.5µL/hour (∼4mg/kg/day) over seven days. Pumps were implanted attached to a catheter connected to a 3.5mm cannula placed into the lateral cerebral ventricle using the following coordinates (with Bregma as landmark): AP-1.3, L 2.0, V 3.5 mm.

### Novel Object Recognition (OR) and Object Location (OL) tests

Tests of spatial memory were conducted four, eight and 12-months following administration of antagonist (Figure 1). Rats were handled and habituated to the testing arena 55×31×21cm for 3 days. For the both procedures, training consisted of rats exploring two identical objects placed 5 cm away from the upper and lower left corners of the testing cage for 10 min. To test 24h recognition memory (ORM) (Figure 2a), rats were presented with a duplicate of a previously encountered object from training and a novel object. For OL memory testing (Figure 2d), one of the two objects was moved to the center of the cage, and the other object remained in the previous location. To exclude any potential location preferences, the familiar locations were counterbalanced within each group. Objects used for both tests were light bulbs, padlocks, yellow radiation containers, large metal clips and beakers, and were used in such a way to avoid repetition between animals. For both OR and OL tests the duration of exploration of each object (sniffing with the animal’s nose < 2 cm from objects) as well as total object exploration time were recorded during the 5 min sessions. The discrimination index (DI) was calculated (novel - familiar)/(novel + familiar))*100 as an index of memory in the ORM and OLM tests.

**Figure 1.**
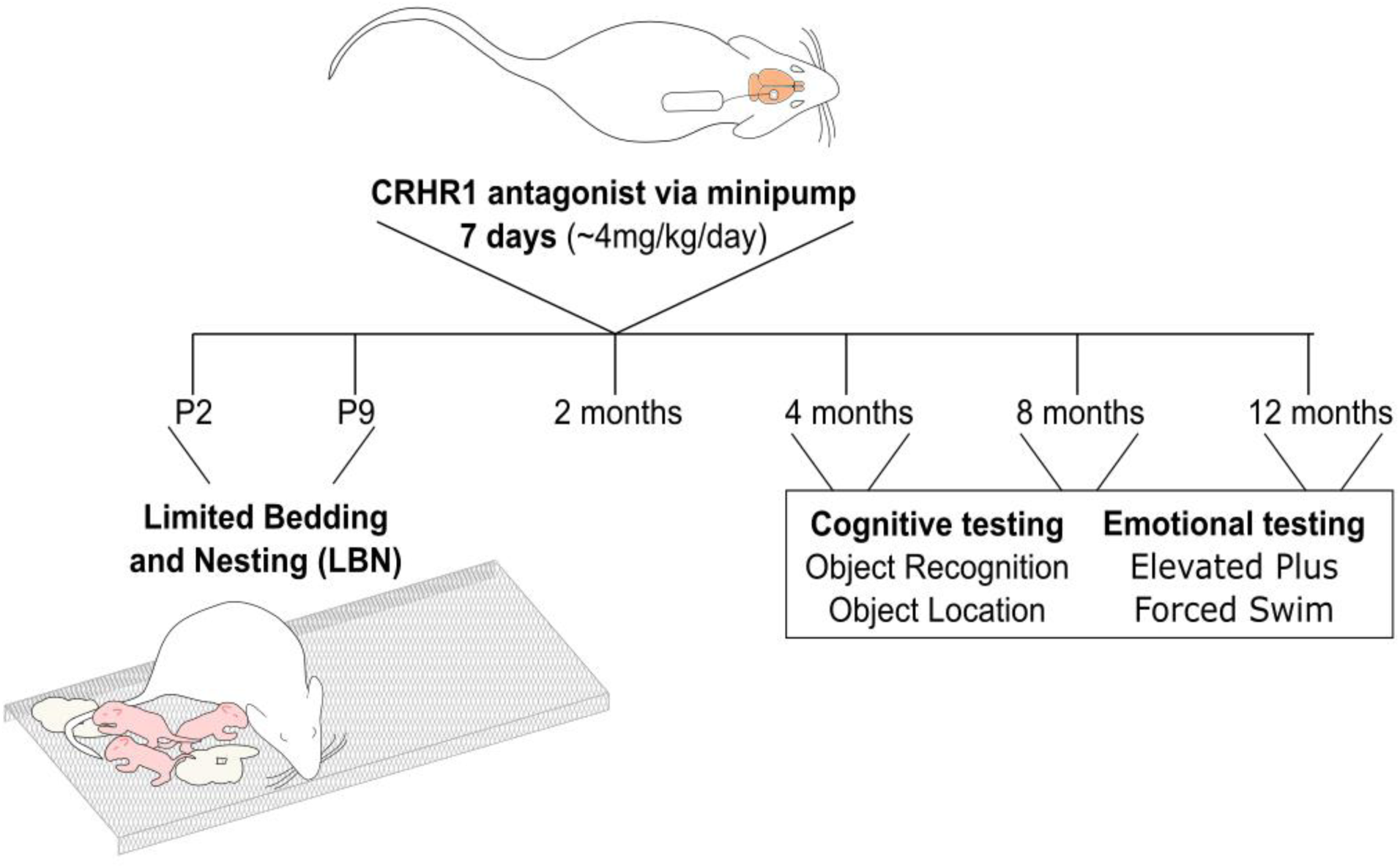
Experimental Design. Experimental timeline. Animals underwent surgery at 2 months of age, a minipump was implanted into the right lateral ventricle and the pump was placed under the skin above the right shoulder. Timeline represents the order of interventions and testing and the ages at which they occurred. LBN = Limited Bedding Nesting, ORM = Object Recognition Memory, OLM = Object Location Memory, EPM = Elevated Plus Maze, OFT = Open Field Test, FST= Forced Swim Test.

**Figure 2.**
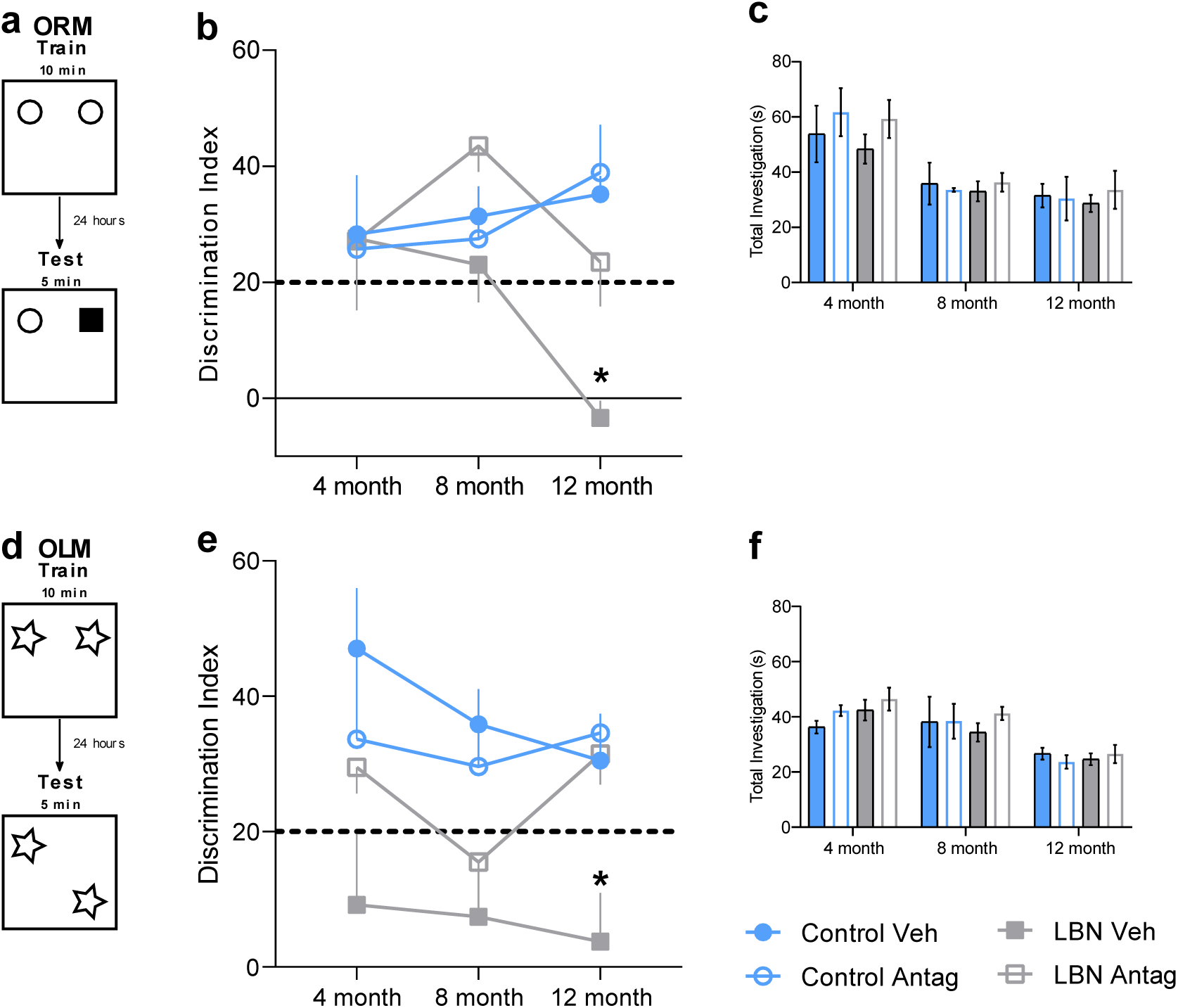
CRHR1 antagonist administration has age dependent effects on the learning and memory impairments following LBN. (A) Animals were trained on the object recognition task (ORM) for 10 minutes, 24 hours later one object was changed, and animals were testing for object discrimination for 5 minutes. (B) There were no effects of CRHR1 antagonist in any of the groups at 4 or 8 months of age, at 12 months of age there was a significant improvement of object discrimination in LBN animals which received administration of CRHR1 antagonist. (C) There were no significant changes in total time the objects were investigated on test day. (D) Animals were trained on new objects for the object location memory (OLM) task, and on testing day, one of these objects was moved to the opposite side of the arena. (E) There was an overall effect of LBN across the ages on the discrimination of the new location, at 12 months of age there was a significant improvement in DI in the LBN animals which received antagonist. (F) There were no differences in the total amount of time the animals spent investigating the objects on test day. Control veh n=3, control antag n=3, LBN veh n=6, LBN antagonist n=6. *p<0.05 (post-test), dots represent mean with ±SEM connected by interpolated lines. ORM = Object Recognition Memory, OLM = Object Location Memory, veh=vehicle, antag=CRHR1 antagonist, LBN = Limited Bedding Nesting.

### Elevated Plus Maze

To examine if the rats exhibited ‘anxiety-like’ behaviors, animals were tested on the elevated plus and open field paradigms. The elevated plus maze consisted of two open arms (50 × 10 cm) and two enclosed arms (50 × 10 × 40 cm) and was elevated 50 cm above the floor. The maze was arranged such that the two arms of each type were opposite each other. Rats were placed in the center of the maze facing an enclosed arm at the start of the experiment. Each rat was allowed one 5-minute trial in the maze, and the maze was cleaned with 70%ethanol after each trial. Decreased time spent on open arms relative to closed arms, compared with control animals, was used as an index of anxiety. Locomotion (the number of times the rat entered the open or closed arms) was also recorded as a measure of activity level.

### Porsolt’s Forced Swim test (FST)

The swim test consists of two sessions separated by 24 hours, in a room illuminated with dim lighting. The habituation session (Day 1), lasts 15 min. Two rats are placed in a glass cylinder (20 cm in diameter and 60 cm high) containing water (25°C) filled to a depth of 45 cm. The test session occurs 24 hours later, and rats are placed in the cylinder for 5 min. Behavior is captured using a video camera, and experimenters monitor rats’ well-being. The durations of floating (immobility), climbing and swimming are scored and serve as indicators of depressive-like vs coping-like behaviors. Water is replaced and containers cleaned between trials.

### Statistical analysis

Three-way repeated measures ANOVAs with a Greenhouse-Geisser correction were used to analyze OLM and ORM data. To account for the missing animal in the 12-month time point, a mixed-effects model (REML) was used to analyze EPM and FST data. Bonferroni correction was used for post-hoc tests to determine specific effects of antagonist. Significance was set at p<0.05. Statistical analyses were performed using GraphPad prism 8.0 (GraphPad software, Inc., LA Jolla, CA). All graphs show the mean ±standard error of the mean (SEM).

## Results

### Following early adversity, CRHR1 antagonist administered during early adulthood rescues learning and memory deficits in middle age

To determine the effects of blocking CRHR1 in adulthood on the LBN-associated deficits in learning and memory, rats were tested on ORM and OLM at four, eight and 12 months of age.

#### Object Recognition Memory (ORM)

No animals had an object preference above 21 on training day; therefore, all animals were included in analysis. There were no overall effects of age (F _(1.79, 25.02)_ = 1.15, p = 0.33), LBN (F _(1, 14)_ = 1.56, p = 0.23), or antagonist (F _(1, 14)_ = 1.34, p = 0.27) on DI during training. No interactions between age x LBN (F _(2, 28)_ = 0.37, p = 0.69) or x antagonist (F _(2, 28)_ = 1.10, p = 0.35), LBN x antagonist (F _(1, 14)_ = 0.10, p = 0.76), or age x LBN x antagonist (F _(2, 28)_ = 1.81, p = 0.18). This suggests no effects of antagonist or LBN on training.

During testing, there were no overall effects of age (F _(1.65, 23.17)_ = 0.75, p =0.46), LBN (F _(1, 14)_ = 2.19, p = 0.16) or antagonist (F _(1, 14)_ = 2.06, p = 0.17) (**Figure 2b**). As expected, there was a significant age x LBN interaction (F _(2, 28)_ = 3.51, p = 0.04), however no age x antagonist (F _(2, 28)_ = 0.87, p = 0.43), LBN x antagonist (F _(1, 14)_ = 2.58, p = 0.13) interactions (**Figure 2b**). Additionally, there was no significant interaction between age x LBN x antagonist (F _(2, 28)_ = 0.47, p = 0.63) (**Figure 2b**). Given the effect of LBN was age dependent, planned post-hoc testing to look at the effect of antagonist revealed no effect of antagonist on controls at four (p>0.05) eight (p>0.05) or 12 (p>0.05) months (**Figure 2b**). CRH antagonist given to animals at two months of age following LBN had no significant effects on object discrimination at four months (p>0.05) or eight months (p>0.05), however there was a significant improvement in object discrimination at 12 months of age (p<0.05) (**Figure 2b**). Therefore, the effect of age-related vulnerability following LBN was inhibited by blocking CRH transiently in adulthood.

Total time investigation the objects was also analyzed as a control measure (**Figure 2c**). There was an overall effect of age on the time investigating the objects during the testing phase (F _(1.96,27.42_= 20.45, p<0.0001) with exploration times decreasing with age (**Figure 2c**). However, there were no effects of LBN (F _(1, 14)_ = 0.10, p = 0.14) or antagonist (F _(1, 14)_ = 0.95, p=0.34) on total object exploration time (**Figure 2c**). No age x LBN (F _(2, 28)_ = 0.15, p = 0.86) or time x antagonist (F _(2, 28)_ = 0.65, p = 0.53) LBN x antagonist (F _(1, 14)_ = 0.36, p = 0.56) or age x LBN x antagonist (F _(2, 28)_ = 0.02, p=0.98) interactions were found, suggesting age effects total investigation time equally between groups (**Figure 2c**).

#### Object Location Memory (OLM)

No animals displayed a significant object preference during training, and all animals were included in analysis. There were no main effects of age (F _(1.92, 26.93)_ = 0.61, p = 0.55) LBN (F _(1, 14_ = 1.40, p = 0.26) or antagonist (F _(1, 14)_ = 1.75, p = 0.21) on DI during training. No age x LBN (F _(2, 28)_ = 0.27, p = 0.77) or x antagonist (F _(2, 28)_ = 0.92, p = 0.41), LBN by antagonist (F _(1, 14)_ = 1.39, p = 0.26), or age x LBN x antagonist (F _(2, 28)_ = 0.62, p = 0.55) interactions were described. Indicating age, LBN or antagonist do not affect training on the OLM task.

During testing for OLM, there was no overall effects of age (F _(1.59, 22.27)_ = 0.92, p = 0.39) or antagonist (F _(1, 14)_ = 3.06, p = 0.10) (**Figure 2e**). As described previously (Ivy et al., 2010; Molet, Maras, et al., 2016), there was an overall effect of LBN on DI’s on test day (F _(1, 14)_ = 24.38, p = 0.0002), indicating a significant impairment of memory at all age groups following LBN (**Figure 2e**). There were no interactions of age with LBN (F _(2, 28)_ = 0.19, p = 0.83) or antagonist (F _(2, 28)_ = 0.96, p = 0.39), nor an age x CES x antagonist interaction (F _(2, 28)_ = 0.35, p = 0.71) (**Figure 2e**). However, LBN significantly determined the effect of the antagonist (F _(1, 14)_ = 9.543, p = 0.008) (**Figure 2e**). Planned post-hoc tests to determine antagonist effects showed no differences in controls at any age group (p>0.05), nor significant differences at 4 or 8 months (p>0.05) in the LBN group. There was, however, a significant rescue of learning and memory at 12 months of age in the group which received the CRHR1 antagonist (p<0.05) (**Figure 2e**). LBN animals were impaired on LBN from four months of age, but CRHR1 antagonist only improved memory at 12 months of age.

As described in ORM, there was a significant effect of age on total exploration time during testing (F _(1.84, 25.82)_ = 20.84, p<0.0001) but no main effects of LBN (F _(1, 14)_ = 0.43, p = 0.52) or antagonist (F _(1, 14)_ = 1.03, p = 0.33) (**Figure 2f**). No age x LBN (F _(2, 28)_ = 0.64, p = 0.53), age x antagonist (F _(2, 28)_ = 0.57, p = 0.57), LBN x antagonist (F _(1, 14)_ = 0.36, p = 0.55) or age x LBN x antagonist (F _(2, 28)_ = 0.37, p = 0.70) were significant (**Figure 2f**).

### Early-life adversity and CRHR1 antagonist administered in early adulthood have no effects on anxiety- and depression-related phenotypes

Given the important relationship of CRH and stress on anxiety- and depression-related phenotypes the animals were tested on the on the elevated plus maze (EPM) and Forced Swim Test (FST).

#### Elevated Plus Maze

When tested for anxiety-like phenotypes on the elevated plus maze there was an overall effect of age on the percentage time spent in the open arm (F _(1.28, 17.33)_ = 4.75, p = 0.04) (**Figure 3a**). There were no main effects of LBN (F (_1, 14)_ = 0.00, p = 0.94) or antagonist (F _(1, 14)_ = 3.96, p = 0.07) (**Figure 3a**). No significant age x LBN (F _(2, 27)_ = 2.41, p = 0.11), age x antagonist (F _(2, 27)_ = 0.18, p = 0.83), LBN x antagonist (F _(1, 14)_ = 0.12, p = 0.74), or age x LBN x antagonist (F _(2, 27)_ = 0.72, 0.05) interactions were reported (**Figure 3a**).

**Figure 3.**
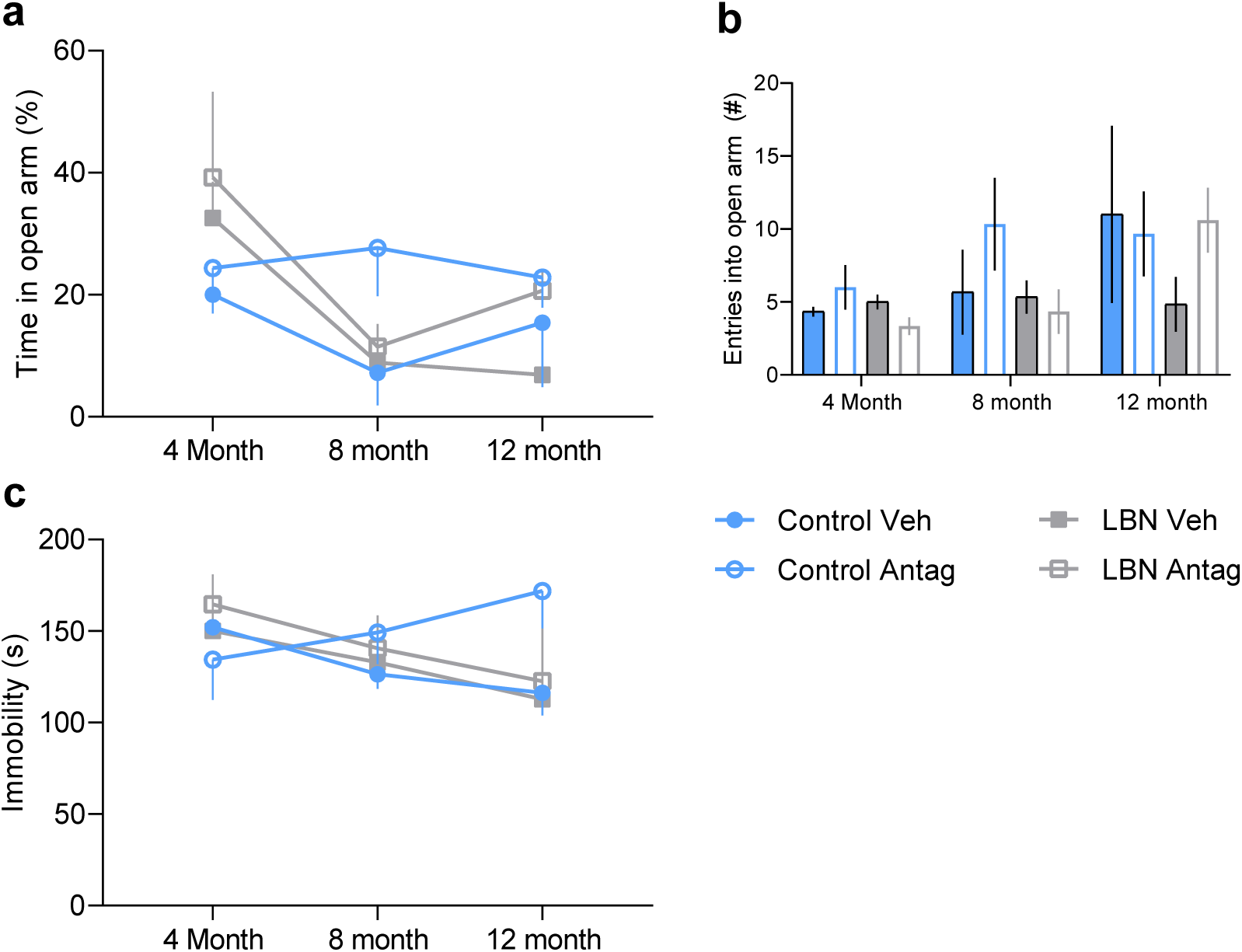
Early-life adversity and CRHR1 antagonist administered to adult rats has no effects on anxiety- and depression-related phenotypes. There was no effect of LBN or CRHR1 antagonist at any of the ages tested on either % time in the open arm (A) or the number of entries into the open arm (B) on the elevated plus maze. (C) Total immobility time in the forced swim test was not affected by LBN nor CRHR1 antagonist. Control veh n=3, control antag n=3, LBN veh n=6, LBN antagonist n=6. *p<0.05 (post-test), dots represent mean with ±SEM connected by interpolated lines. ORM = Object Recognition Memory, OLM = Object Location Memory, veh=vehicle, antag=CRHR1 antagonist, LBN = Limited Bedding Nesting.

Similarly, there was an overall effect of age on the number of entries into the open arm of the maze (F _(1.36, 27.94_) = 4.239, p = 0.04) with no main effects of LBN (F (_1, 41)_ = 3.41, p = 0.07), antagonist (F _(1, 41)_ = 1.21, p = 0.28) (**Figure 3b**). There were also no significant interactions of age x LBN (F _(2, 41)_ = 0.28, p = 0.75), age x antagonist (F _(2, 41)_ = 0.31, p = 0.73), LBN x antagonist (F _(1, 41)_ = 0.07, p = 0.80), nor age x LBN x antagonist (F _(2, 41)_ = 2.54, p = 0.09) (**Figure 3b**). While, age or repeated testing decreases EPM performance, it is consistent between the groups.

#### Forced Swim Test

There were no main effects of age (F _(1.28, 17.28)_ = 1.82, p = 0.19), LBN (F _(1, 14)_ = 0.05, p = 0.83) or antagonist (F _(1, 14)_ = 0.84, p = 0.38) or total time immobile in the forced swim test (**Figure 3c**). Additionally, there were no age x LBN (F _(2, 27)_ = 1.90, p = 0.17), age x antagonist (F _(2, 27)_ = 1.65, p = 0.21), LBN x antagonist (F _(1, 14)_ = 0.06, p =0.82) or age x LBN x antagonist (F _(2, 27)_ = 1.79, p = 0.19) interactions (**Figure 3c**). Suggesting no changes to depression-related phenotypes in any of the groups on the FST.

## Discussion

These experiments indicate that interventions, even when administered outside of the sensitive period, may ameliorate deficits observed in learning and memory following early-life adversity (Bath et al., 2016; Brunson et al., 2005; Molet, Maras, et al., 2016; Naninck et al., 2015; Rice et al., 2008). We have previously demonstrated that when a CRHR1 antagonist is administered directly following a period of early-life adversity, within sensitive developmental periods, there are significant improvements in learning and memory as early as two months of age (Ivy et al., 2010). However, here we show improvements on ORM and OLM only at 12 months of age, 10 months after the CRH antagonist was administered.

Previous work has indicated that the effects of LBN on learning and memory become more prominent when brain circuits are challenged (Brunson et al., 2005; Ivy et al., 2010; Molet, Maras, et al., 2016). The ability to discriminate new objects in the testing phase of ORM utilizes multiple brain regions, including the hippocampus and the perirhinal cortex (Winters, Saksida, & Bussey, 2008), whereas discrimination in OLM is considered largely hippocampal dependent (Mumby, Gaskin, Glenn, Schramek, & Lehmann, 2002). Previous work has described decreased learning and memory on OLM following early-life adversity as early as two months, while the ability to perform the ORM task is intact until 12 months of age (Ivy et al., 2010; Molet, Maras, et al., 2016). Additionally, following adversity early in life, an animal is more susceptible to memory disruption after a second stress when compared to those with optimal early experiences (Molet, Maras, et al., 2016). This vulnerability to challenges is also observed with ageing (Brunson et al., 2005). While some impairment in hippocampal related memory with ageing is expected (Burke & Barnes, 2006; Morrison & Baxter, 2012), an experience of early adversity results in an acceleration of these deficits (Brunson et al., 2005).

In the present study, the age induced impairments on learning and memory tasks following LBN were rescued by blocking CRHR1 in the adult brain (Figure 2). The age-induced deficit on ORM that occurs in LBN animals at 12 months of age was rescued by administration of the CRHR1 antagonist (Figure 2b). However, where we see a vulnerability to LBN on the OLM tasks as early as 4 months, there is no rescue until 12 months of age (Figure 2e). Previous work has demonstrated that when the antagonist is administered at P10, improvements on the OLM task occur as early as 2 months of age (Ivy et al., 2010).

The developing hippocampus is more sensitive to fluctuations in CRH compared to the adult brain (Brunson, Chen, Avishai-Eliner, & Baram, 2003; Hollrigel, Chen, Baram, & Soltesz, 1998; Maras & Baram, 2012; Smith & Dudek, 1994). Therefore, it is unsurprising that blocking CRHR1 at P10 can cause such staggering improvements in learning and memory. At two months of age, the effects of acute increases in CRH on the brain are more transient, and do not influence hippocampal development (Brunson et al., 2003; Smith & Dudek, 1994). However, the adult hippocampus is responsive to changes in CRH levels and prolonged exposure to CRH reduces synaptic transmission and impairs synaptic plasticity through the loss of spines (Chen et al., 2008, 2013). At 2 months of age CRH levels are increased in LBN animals (Fenoglio et al., 2006; Ivy et al., 2010; Maras & Baram, 2012), therefore blocking CRHR1 may inhibit this CRH induced spine loss and restore synaptic activity within the hippocampus.

While CRH is important for developmental plasticity within the hippocampus (Curran, Sandman, Poggi Davis, Glynn, & Baram, 2017; Dubé et al., 2015; Ivy et al., 2010; Sandman et al., 2018; Singh-Taylor et al., 2018), its role in affective-related disorders is less clear (Spierling & Zorrilla, 2017). This is observed in the present study with a lack of an effect of antagonist on the anxiety- or depression-related phenotypes tested. Impairments in learning and memory following LBN have been well replicated (Bath et al., 2016; Brunson et al., 2005; Molet, Maras, et al., 2016; Naninck et al., 2015; Rice et al., 2008), however, its influence on affective-related disorders is variable. As described in the present study (Figure 3.), previous studies have found no effect on EPM or FST (Bolton et al., 2018; Brunson et al., 2005; Molet, Heins, et al., 2016; Naninck et al., 2015). However, increased anxiety-like phenotypes (Dalle Molle et al., 2012; Guadagno, Wong, & Walker, 2018; Wang et al., 2012) and increased immobility time during FST (Raineki, Cortés, Belnoue, & Sullivan, 2012) have also been described following LBN.

In rodents, early adversity causes memory deficits which worsen with age, likely due to altered learning and memory pathway development. These results suggest that it is possible to reduce impairments in learning and memory by intervening in early adulthood, however they are less effective when given outside of the sensitive period.

